# Potential function of *CbuSPL* and gene encoding its interacting protein during flowering in *Catalpa bungei*

**DOI:** 10.1101/803122

**Authors:** Zhi Wang, Tianqing Zhu, Erqin Fan, Nan Lu, Fangqun Ouyang, Nan Wang, Guijuan Yang, Lisheng Kong, Guanzheng Qu, Shougong Zhang, Wenjun Ma, Junhui Wang

**Author notes:** To whom correspondence should be addressed. (J.H.W); (W.J.M).

## Abstract

“Bairihua”, a variety of the Catalpa bungei, has a large amount of flowers and a long flowering period which make it an excellent material for flowering researches in trees. SPL is one of the hub genes that regulate both flowering transition and development. Here, a SPL homologues *CbuSPL9* was cloned using degenerate primers with RACE. Expression studies during flowering transition in Bairihua and ectopic expression in Arabidopsis showed that *CbuSPL9* was functional similarly with its Arabidopsis homologues. In the next step, we used Y2H to identify the proteins that could interact with CbuSPL9. HMGA, an architectural transcriptional factor, was identified and cloned for further research. BiFC and BLI showed that CbuSPL9 could form a heterodimer with CbuHMGA in the nucleus. The expression analysis showed that *CbuHMGA* had a similar expression trend to that of *CbuSPL9* during flowering in “Bairihua”. Intriguingly, ectopic expression of *CbuHMGA* in Arabidopsis would lead to aberrant flowers, but did not effect flowering time. Taken together, our results implied a novel pathway that *ChuSPL9* regulated flowering development, but not flowering transition, with the participation of *ChuHMGA*. Further investments need to be done to verify the details of this pathway.

## Introduction

Flowers allow flowering plants to have a broader evolutionary relationship and extend their ecological niche so that they can dominate the terrestrial ecosystem. Flowering is extremely important for the development of perennial woody plants and for improving the economic value of plants. However, due to complex genomes and other objective characteristics of perennial woody plants, research on the flowering process in perennial woody plants remains limited. *Catalpa bungei* is valuable as both a timber and an ornamental tree^1^. “Bairihua”, which is a natural variety of *C. bungei*, has been characterized for its especially short juvenile period, large number of flowers and long flowering period. The flowering period of “Bairihua” is approximately 15 days, and its accumulative flowering period reaches 100 days, which is very rare for woody plants (http://www.forestry.gov.cn/). “Bairihua” provides an excellent opportunity to evaluate the flowering process of woody plants.

Flowering is controlled by sophisticated regulatory networks^2–5^. Five major pathways are involved in these processes, including the aging pathway^6^, gibberellin pathway^7–11^, photoperiod pathway^12–16^, vernalization pathway^17–19^ and autonomous pathway^20^. The SQUAMOSA promoter-binding protein-LIKE (SPL) family of transcription factors (TFs) integrate multiple pathways^21–25^. *SPLs* have been shown to regulate flowering time and flower organ development in both herbs and woody plants, such as *Gossypium hirsutum*^26^, maize^27^, birch^28^, *Prunus mume*^29^, and *Platanus acerifolia*^30^. In the model plant Arabidopsis, AtSPLs have been shown to be a group of dominant regulators of the flowering process^4, 11, 13, 22, 24, 31–34^. The overexpression of *AtSPLs* leads to early flowering and abnormal inflorescence, and conversely, the inhibition of *AtSPL* expression delays the occurrence of floral transition^21, 35–37^. As a group of TFs, SPLs regulate the expression of other genes. Numerous downstream genes of SPLs have been identified; for example, *AtSPL3* can directly upregulate the expression of *LFY*, *FUL* and *AP1* by binding to their promoters^38–40^. However, in addition to protein-DNA interactions, TFs also affect plant growth and development by forming protein complexes. For example, the MYB-bHLH-WD40/WDR (MBW) complex regulates late biosynthetic genes in anthocyanin biosynthesis, impacts fruit quality in apple^41, 42^, and regulates trichome initiation in *Arabidopsis thaliana*^43^. Two TFs, AT-HOOK MOTIF NUCLEAR LOCALIZED PROTEIN 3/4, regulate the formation of the tissue boundary between the procambium and xylem in Arabidopsis roots^44^. Few studies have investigated whether there are other factors that interact with SPLs and affect their binding ability, especially in trees. The function of SPLs in woody plants is still in its infancy.

In an effort to study the molecular mechanism of the flowering process in “Bairihua”, we evaluated whether there were protein interactions involved in SPL regulation during the flowering process. As a first step to address this question, we isolated and characterized the *SPL9* orthologous gene from “Bairihua” and performed native in planta gene expression analysis. The putative function of this gene was then tested by ectopic expression experiments. This gene showed similar expression patterns and the ability to induce flower organ development and early flowers in Arabidopsis. The findings of this study indicated that *CbuSPL9* is functionally conserved. An architectural TF, CbuHMGA, was found via screening CbuSPL9-interacting proteins. *CbuHMGA* is involved in the floral organ development of “Bairihua”’, but not in the regulation of flowering time. These results provide a molecular basis for studying the molecular mechanism of “Bairihua” flowering and provide a research direction for the study of the floral transition of perennials.

## Materials and methods

### Plant materials

*C. bungei* is a perennial tree that typically flowers over a 30-day flowering period. However, “Bairihua”, the new variety of *C. bungei*, was found in Henan Province, China, and confirmed by the National Forestry and Grassland Administration (http://www.forestry.gov.cn/). The flowering period of a single flower is approximately 15 days, and its accumulative flowering period reaches 100 days, which is very rare for woody plants. From January 15 to April 2, 2017, we collected the first round of axillary buds of EF and NF varieties every one to two days. Samples were collected every 10 days during the dormant and germinating periods. Since the floral transition of “Bairihua” was completed within 7 to 10 days, samples were collected every day during the floral transition period, and the samples were collected every 5 days during the reproductive growth stage. The samples used for RNA extraction were washed with distilled water, frozen immediately in liquid nitrogen, and stored at −80°C. Samples for histological analysis were fixed in a formalin:glacial acetic acid:70% ethanol (5:5:90 vol.; FAA) solution under a vacuum for at least 24 h.

### Histological analysis

For histological analysis, the samples were immersed in FAA fixative and placed under vacuum at 4°C. Samples were dehydrated in gradient ethanol and then embedded in paraffin. Ten-mm-thick sections (RM2255 Fully Automated Rotary Microtome; Leica, Germany) were stained with Safranine O and fast green FCF (Sigma-Aldrich, USA). The slices were observed and photographed using a Leica DM 6000B fully automated upright microscope (Leica Microsystems GmbH, Wetzlar, Germany).

### Cloning *CbuSPL9* and *CbuHMGA* sequences from “Bairihua”

RNA from “Bairihua” was extracted from the buds of the mutant using an RNA extraction kit (TaKaRa), and contaminating DNA was removed with RNase-free DNase I (TaKaRa). One or two micrograms of total mRNA template was added to Oligod (dT)18 primer and reverse-transcribed into single-stranded cDNA by M-MLV RTase (TaKaRa). The full-length cDNA of *CbuSPL9* was cloned via 3’-RACE and 5’-RACE by using the (TaKaRa) according to the manufacturer’s instructions. In 3’-RACE, the *CbuSPL9* gene-specific forward primers F1/F2 were designed based on published and aligned *SPL* sequences from NCBI (http://blast.ncbi.nlm.nih.gov/Blast.cgi). F1 nested immediately upstream of F2. The 3’-cDNA synthesis primer was provided in a kit. The PCR products were cloned into the PMD18-T vector and sequenced. *CbuSPL9s* were identified using BLAST. In 5’-RACE, the gene-specific reverse primers R1/R2 (Supplementary Table S1) were designed based on sequences from 3’-RACE. R2 nested immediately upstream of R1. The PCR products were cloned into the PMD18-T vector and sequenced.

### Yeast two-hybrid (Y2H) screening of the “Bairihua” cDNA library

A yeast library was constructed by Oebiotech (Shanghai, China) via cloning the full-length cDNA library from mRNAs in the “Bairihua” buds into the pGADT7 vector^46^. *CbuSPL9* was inserted into the pGBKT7 vector. Screening of the yeast library was performed using pGBKT7-*CbuSPL9*. The corresponding primers are listed in Supplementary Table S1. Transformed cells were grown on SD medium supplemented with Trp. The transformants were screened on supplemented SD medium lacking Leu, Trp, His and Ade and supplemented with X-a-Gal and Aureobasidin A. Plates were incubated at 30°C for 48-72 h and photographed. The positive clones were verified with both pGBKT7 vector consensus primers. Positive control mating was as follows: pGADT7-T in Y2HGold and pGBK-53 in Y187. Negative control mating was as follows: pGADT7-T in Y2HGold and pGBKT7-Lam in Y187.

### Subcellular localization

To validate subcellular localization, the full-length coding sequences (without the stop codon) of *CbuSPL9* and *CbuHMGA* were amplified from RNA of “Bairihua” buds by RT-PCR. The PCR products of *CbuSPL9* and *CbuHMGA* were ligated to the vector pCAMBIA1304 using the Seamless Assembly Cloning Kit (CloneSmarter, Beijing, China) to construct the CbuSPL9/CbuHMGA-GFP fusion genes driven by a CaMV35S promoter (Niwa, 2003). pCAMBIA1304-GFP was used as a positive control. The transient expression vectors *CbuSPL9*-GFP, *CbuHMGA*-GFP and GFP-HDEL were injected into the leaf lower epidermal cells of *Nicotiana tabacum* L. using Agrobacterium transformation as described by Chen et al. (2011). The transformed cells were incubated for 2 days. The leaves were removed, cut into squares and immersed in PBS buffer containing 1 g mL^-1^ DAPI to stain the nuclei. The transient expression of the CbuSPL9/CbuHMGA-GFP fusion proteins was observed under an UltraVIEW VoX 3D Live Cell Imaging System Spinning Disk confocal laser scanning microscope (PerkinElmer, Waltham, MA, USA). The wavelength of excitation used was 488 nm for GFP and 405 nm for DAPI.

### BiFC analysis

To confirm and visualize the interaction between CbuSPL9 and CbuHMGA in protoplasts from *Populus trichocarpa*, a BiFC assay was performed based on split EYFP. EYFP was fused to the C-terminus of CbuSPL9 and the N-terminus of CbuHMGA, resulting in CbuSPL:EYFP^C^ and CbuHMGA:EYFP^N^. A positive EYFP signal indicates an interaction of EYFP^C^ and EYFP^N^ due to the heterodimerization of CbuSPL9 with CbuHMGA. CbuSPL:EYFP^C^ was cotransfected with CbuHMGA:EYFP^N^ and H2A:mCherry into protoplasts. The transient expression of the CbuSPL9/CbuHMGA-GFP fusion proteins was observed under an UltraVIEW VoX 3D Live Cell Imaging System Spinning Disk confocal laser scanning microscope (PerkinElmer, Waltham, MA, USA).

### Biolayer interferometry assay

CbuSPL9 was cloned into pGEX6P-1 as a C-terminal GST-tagged construct, and the construct was confirmed by sequencing. CbuHMGA was cloned into pET28a as an N-terminal 6His-tagged construct, and the construct was confirmed by sequencing. The protein was purified using the procedure for EMCV-3C and RV-3C as described above. Real-time interactions between CbuHMGA and CbuSPL9 were monitored with an Octet QK (Forte-Bio) that is based on BLI^47^. BLI was used to determine dissociation constants (K_D_) as well as the on-and off-rate (k_on_ and k_off_) for HIS-CbuHMG binding to GST-CbuSPL1.

### Bioinformatic analysis

Multiple sequence alignment and phylogenetic analysis were performed using MEGA6.0. After alignment, the evolutionary history was calculated using the neighbor-joining (NJ) method. The tree was inferred from 1000 bootstrap replicates to show the evolutionary history of the genes. The MEME online tool (http://meme-suite.org/tools/meme) was used to identify the motifs of the CbuSPL9 protein. MEME was run locally with the following parameters: number of repetitions = any and maximum number of motifs = 20. All candidate interacting protein sequences were examined by the domain analysis program SMART (Simple Modular Architecture Research Tool) (http://smart.embl-heidelberg.de/).

### RNA extraction and quantitative real-time PCR

Total RNA was isolated from “Bairihua” buds at different developmental stages. The purity and quality of RNA was checked by NanoDrop8000 (Thermo Fisher Scientific, Waltham, MA, USA) and analyzed by gel electrophoresis. First-strand cDNA synthesis was carried out with ∼1 μg RNA using the SuperScript III reverse transcription kit (Invitrogen) and random primers according to the manufacturer’s instructions. Primers were designed using Primer 3 online. The melting temperature of the primers was 60°C, and the amplicon lengths were 100-200 bp. All primers are listed in Supplementary Table S1. qRT-PCR was performed on a 7500 Real-Time PCR System (Applied Biosystems, CA, USA) using a SYBR Premix Ex Taq™ Kit (TaKaRa, Dalian, China) according to the manufacturer’s instructions. Relative expression levels were calculated using the 2-ΔΔCt method. Actin was used as an internal control, and each reaction was conducted in triplicate. The stem expression values were set to 1.

### Transient overexpression in *Arabidopsis thaliana*

Full-length CbuSPL9 and CbuHMGA were cloned into the binary vector pBI121 (BD Biosciences Clontech, USA) under the control of the cauliflower mosaic virus 35S promoter in the sense orientation. The transgenic plants were generated with the 35S:CbuSPL9 and 35S:CbuHMGA constructs via *Agrobacterium tumefaciens* GV3101 by using the floral dip method (Clough and Bent, 1998). Surface-sterilized T1 seeds were grown on a solid 0.5 × MS medium containing 30 μg mL^-1^ hygromycin at 4°C for 2 days, which were then transferred to the greenhouse under long-day conditions (16 h light/8 h dark) at 22°C for 10 days. Subsequently, the seedlings were transplanted into soil. Phenotypes of the transgenic plants were observed in the T1 generation, and the overexpression of *CbuSPL9* and *CbuHMGA* in the transgenic plants was confirmed by PCR genotyping (Supplementary Figure S1 and S2). For each construct, at least 10 transgenic lines with similar phenotypes were observed, and 3 of them were used for detailed analysis.

### Flowering time measurement

Flowering time was measured by counting the total number of rosette leaves and the number of days to flower (when the floral buds were visible). The numbers of rosette and cauline leaves of ∼20 plants were counted and averaged. The presence of abaxial trichomes was used to differentiate between the juvenile and adult vegetative leaves. Data were classified with Win-Excel and analyzed via analysis of variance (ANOVA) using the SPSS (version 8.0, SPSS Inc., Chicago, IL, USA) statistical package. Comparisons between the treatment means were made using Tukey’s test at a probability level of P <=0.05.

## Result

### A flowering-related *SPL* homologous gene was isolated in “Bairihua”

The SPL homologous genes in *C. bungei* were isolated by using degenerate primers targeting the homeodomain. The isolated homeodomain sequences were extended by genome walking to acquire the full genomic sequences (exons and introns). One of the isolated sequences encoded a protein with a typical SPL protein structure, which included a highly conserved SBP-box domain bearing two zinc-binding sites and one bipartite nuclear localization signal^45^ (Supplementary Table S2). The first Zn-finger-like structure (ZN-1 in Figure 1a) was C3H-type, and the second (Zn-2 in Figure 1a) was C2HC-type. The nuclear localization sequence (NLS) is a highly conserved bipartite domain located at the C-terminus of SBP. Phylogenetic analysis showed that the isolated sequence clustered with AtSPL9 and AtSPL15 (Figure 1b). Since it shared more sequence similarity with *AtSPL9*, we named the *SPL* homologous gene *CbuSPL9*.

**Figure 1.**
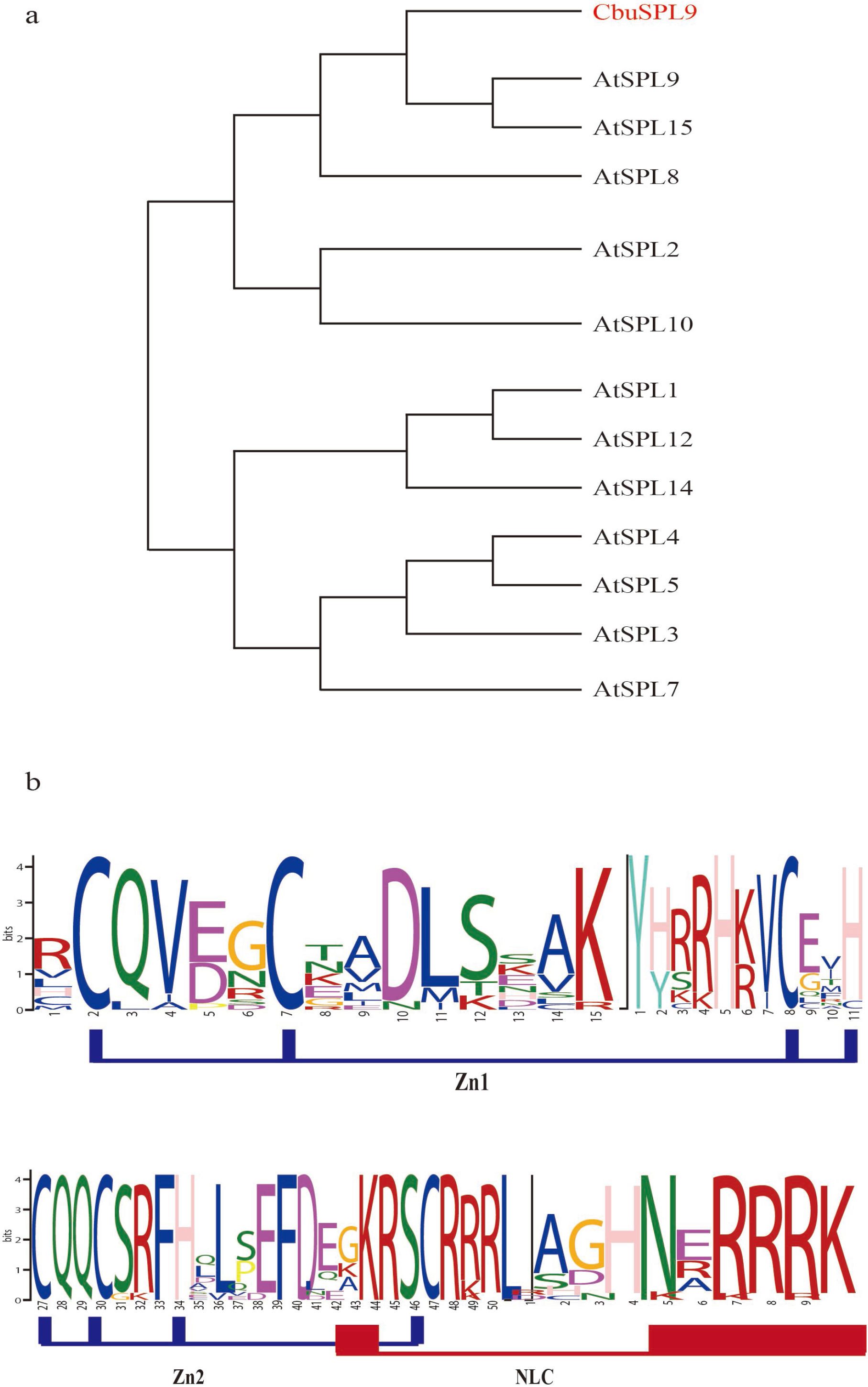
The phylogenetic relationship and motif composition analysis of the CbuSPL9 gene in *C. bungei*. a) Multiple sequence alignment and sequence logo of the *C. bungei* SBP-box domain. Sequence alignment was performed with DNAMAN. The two conserved zinc fingers and NLS are indicated. The sequence logo was obtained from MEME online software. The overall height of the stack indicates the sequence conservation at that position. b) The phylogenetic tree was constructed with MEGA 6.0 by the neighbor-joining (NJ) method with 1000 bootstrap replicates. Bootstrap support is indicated at each node. *A. thaliana* (At), *C. bungei* (Cbu).

### Expression of *CbuSPL9* during the flowering process

Intensive sampling was performed to investigate the changes in *CbuSPL9* expression during the flowering process (Figure 2). T1-T3 was the dormant period during which no significant differences in morphology could be observed between the flowering buds (FBs) and the leaf buds (LBs). T4-T5 was the germination period. During this period, the internal morphology was similar between the FBs and LBs, but the external morphology of the FBs and LBs had a pale green appearance compared with that of the FBs and LBs during the dormant period. T6-T9 followed the short germination period and was the floral transition period, during which flower primordium and leaf primordium were developed in the FBs and LBs, respectively. Finally, T10-T12 was the reproductive growth period. The expression level of *CbuSPL9* was significantly higher in the FBs than in the LBs, especially during the dormant period and the reproductive growth period. Overall, the expression study supported that *CbuSPL9* was an SPL homologous gene and involved in flowering regulation.

**Figure 2.**
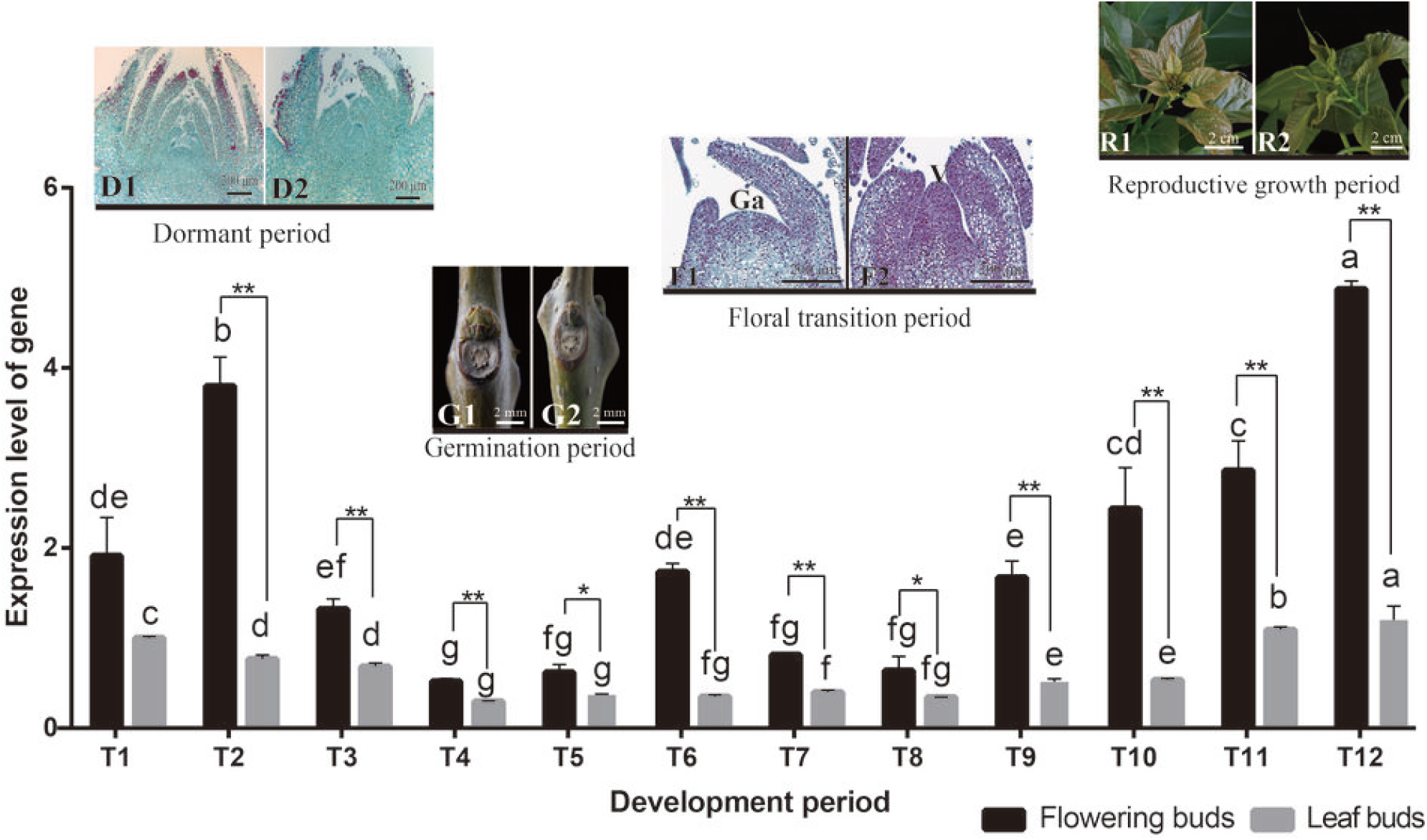
Expression profile of *CbuSPL9* in the flower buds and leaf buds during the developmental periods of *C. bungei*. T1-T3 represents the flowering buds and leaf buds collected in the dormant period; T4-T5 represents the flowering buds and leaf buds collected in the germination period; T6-T9 represents the flowering buds and leaf buds collected in the floral transition period; and T10-T12 represents the flowering buds and leaf buds collected in the reproductive period. D1: Image of flowering buds in the dormant period; D2: Image of leaf buds in the dormant period; G1: Image of flowering buds in the germination period; G2: Image of leaf buds in the germination period; F1: Section of flowering buds in the floral transition period; note the flat generative apex (Ga); F2: Section of leaf buds in floral transition period, note the bulged vegetative apex (V); R1: Image of flowering buds in the reproductive period; and R2: Image of leaf buds in the reproductive period. Notably, even though floral transition was never observed in the leaf buds, the leaf buds collected in the period corresponding to floral transition are henceforth called F2 and R2 for convenience. Error bars indicate SD from three independent biological replicates. *Difference between flowering buds (black) and leaf buds (gray) is significant (Student’s test; p<0.05). **Difference between flowering buds (black) and leaf buds (gray) is highly significant (Student’s test; p<0.01).

### The overexpression of *CbuSPL9* in Arabidopsis

As there is no available transformation system in *C. bungei*, *CbuSPL9* was overexpressed in Arabidopsis (Columbia ecotype, col). The flowers from *col* were tetradynamous and had four petals distributed in cross type (Figure 3aI). In contrast, the *oe-spl9* transgenic plants showed aberrant flower organs (Supplementary Table S3). The *oe-spl9* transgenic lines exhibited flower organs with altered numbers and locations, such as shrunken petals, increased stamens, and overlapping petals (Figure 3a II-VI). In addition to the evident changes in floral organ morphology, an acceleration in flowering time was observed in the *oe-spl9* lines (Figure 3b, Supplementary Table S4). The *col* line initiated flowering when 14 rosette leaves were present (Figure 3c). However, *oe-spl9* possessed less than 9 rosette leaves at the time of bolting (Figure 3d). This result indicated that the function of *CbuSPL9* was conserved. The regulatory mechanism of *CbuSPL9* in “Bairihua” might be similar to that in Arabidopsis.

**Figure 3.**
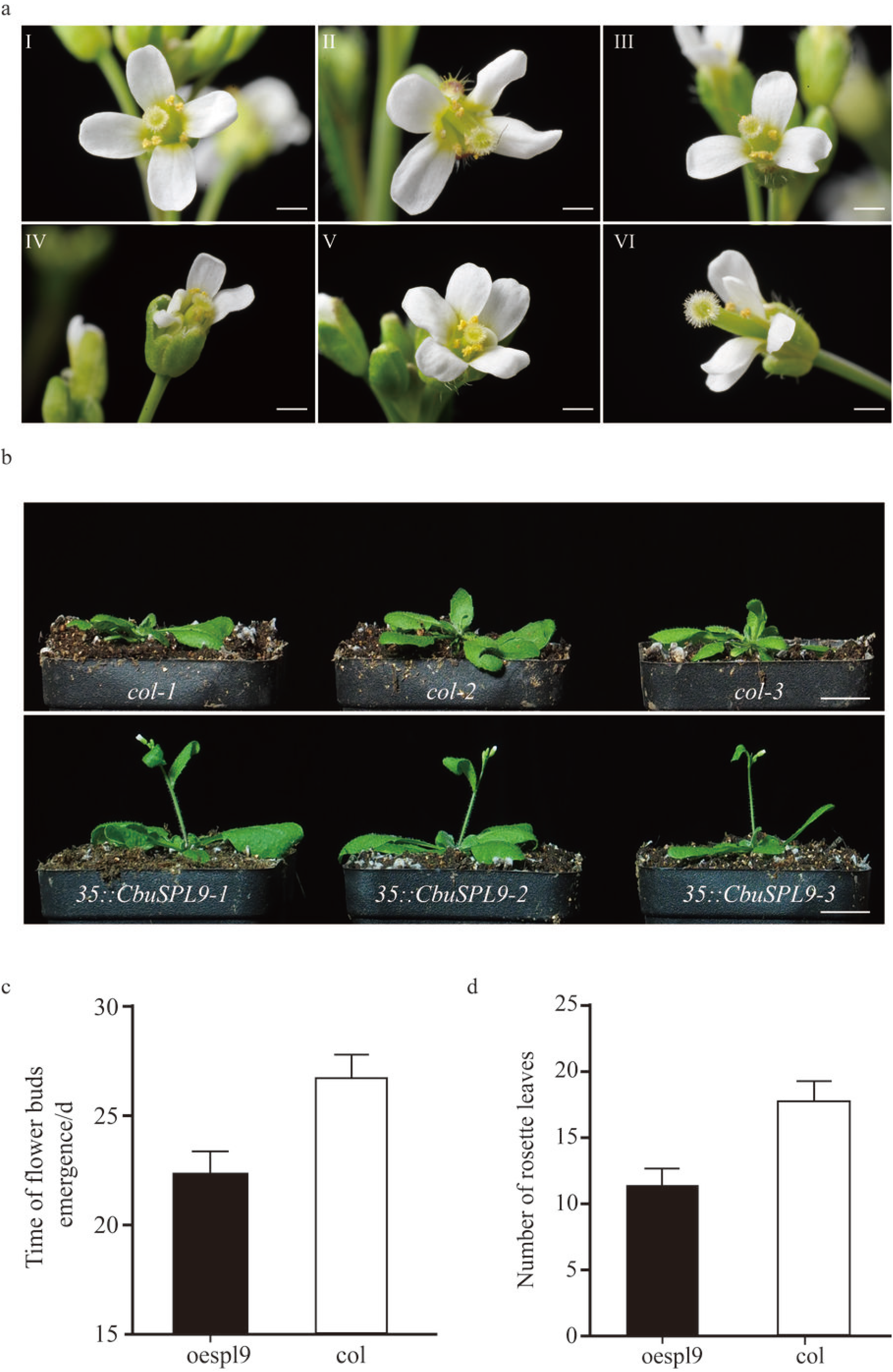
Phenotype of overexpression mutants of *CbuSPL9*. a) The change in floral organs occurred in the CbuSPL9-overexpressing transgenic Arabidopsis lines. Image of a normal flower from col as the control (a/I); and the floral organ mutant from oespl9 (a/II-VI). b) Comparison of the flowering phenotype in both the col and oespl9 transgenic plants. From top to bottom: col (control) and oespl9 (transgenic plants). c) and d) are the statistical data of the time of flower bud emergence and the number of rosette leaves. The oespl9 transgenic plants (black), the oe-hmga transgenic plants (gray) and the col plants (white). A total of 30 plants were averaged to obtain the mean. Error bars indicate SD.

### Screening *CbuSPL9*-interacting proteins

As a TF, SPL-DNA interaction studies have been extensively performed. However, TFs can also fine-tune specific biological processes through protein interactions. We constructed a *C. bungei* yeast two-hybrid cDNA library to explore the proteins that interact with *CbuSPL9*. A total of 809 blue colonies representing potential positive clones were obtained on QDO/Aba/X-a-Gal plates. The potential positive clones were subsequently tested by PCR for the library plasmid, and 406 clones were positive. The resulting PCR products were sequenced, and the sequences were aligned using the NCBI BLASTp search function. Finally, 12 interacting protein candidates were identified (Table 1). These predicted proteins included HMGA, aquaporins, bHLH48, GATA-related protein, PHD finger protein ALFIN-LIKE 4, heavy metal-associated protein, etc.

**Table 1.**
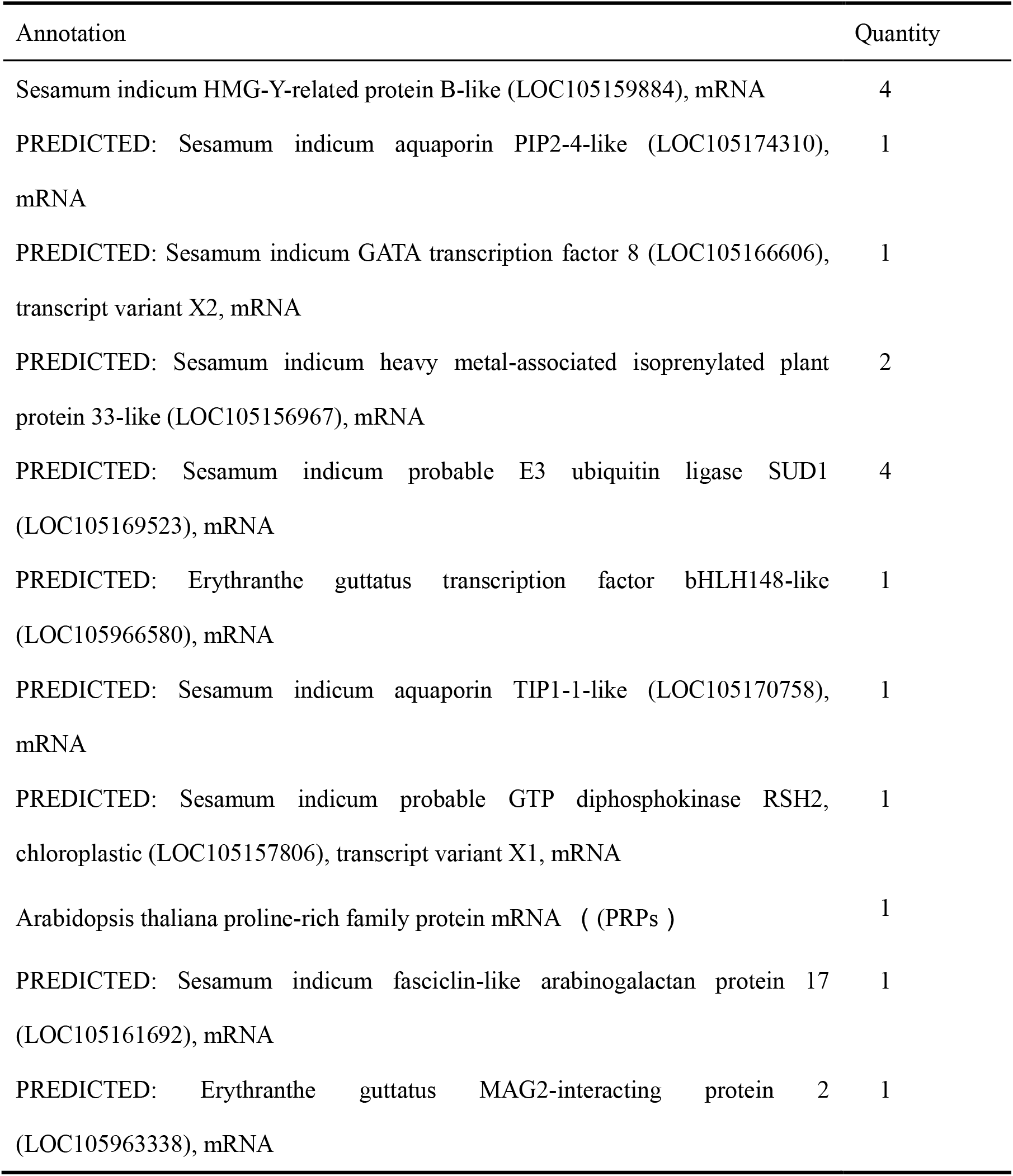
Details of the proteins interacting with the CbuSPL9 protein

| Annotation | Quantity |
| --- | --- |
| Sesamum indicum HMG-Y-related protein B-like (LOC105159884), mRNA | 4 |
| PREDICTED: Sesamum indicum aquaporin PIP2-4-like (LOC105174310), mRNA | 1 |
| PREDICTED: Sesamum indicum GATA transcription factor 8 (LOC105166606), transcript variant X2, mRNA | 1 |
| PREDICTED: Sesamum indicum heavy metal-associated isoprenylated plant protein 33-like (LOC105156967), mRNA | 2 |
| PREDICTED: Sesamum indicum probable E3 ubiquitin ligase SUD1 (LOC105169523), mRNA | 4 |
| PREDICTED: Erythranthe guttatus transcription factor bHLH148-like (LOC105966580), mRNA | 1 |
| PREDICTED: Sesamum indicum aquaporin TIP1-1-like (LOC105170758), mRNA | 1 |
| PREDICTED: Sesamum indicum probable GTP diphosphokinase RSH2, chloroplastic (LOC105157806), transcript variant X1, mRNA | 1 |
| Arabidopsis thaliana proline-rich family protein mRNA ( PRPs ) | 1 |
| PREDICTED: Sesamum indicum fasciclin-like arabinogalactan protein 17 (LOC105161692), mRNA | 1 |
| PREDICTED: Erythranthe guttatus MAG2-interacting protein 2 (LOC105963338), mRNA | 1 |

### Cloning the *HMGA* gene

Among these candidates, HMGA, which is an AT-hook rich protein, belongs to the most abundant nonhistone protein family in the nucleus. We cloned the full-length *HMGA* homologous gene via rapid amplification of cDNA ends. Phylogenic analysis revealed that this protein has a close relationship with AtHMGA. Therefore, this protein was renamed CbuHMGA. *CbuHMGA* encoded a H15 domain and 4 AT-hook motifs (Figure 4a). The expression analysis showed that the expression trend of *CbuHMGA* was similar to that of *CbuSPL9* in the four periods in “Bairihua” (Figure 4b). The expression of *CbuHMGA* was elevated in the dormant period, and the highest expression level was detected in the reproductive growth period. However, the increase in the expression of *CbuHMGA* before the reproductive growth period occurred slightly earlier than that of *CbuSPL9*.

**Figure 4.**
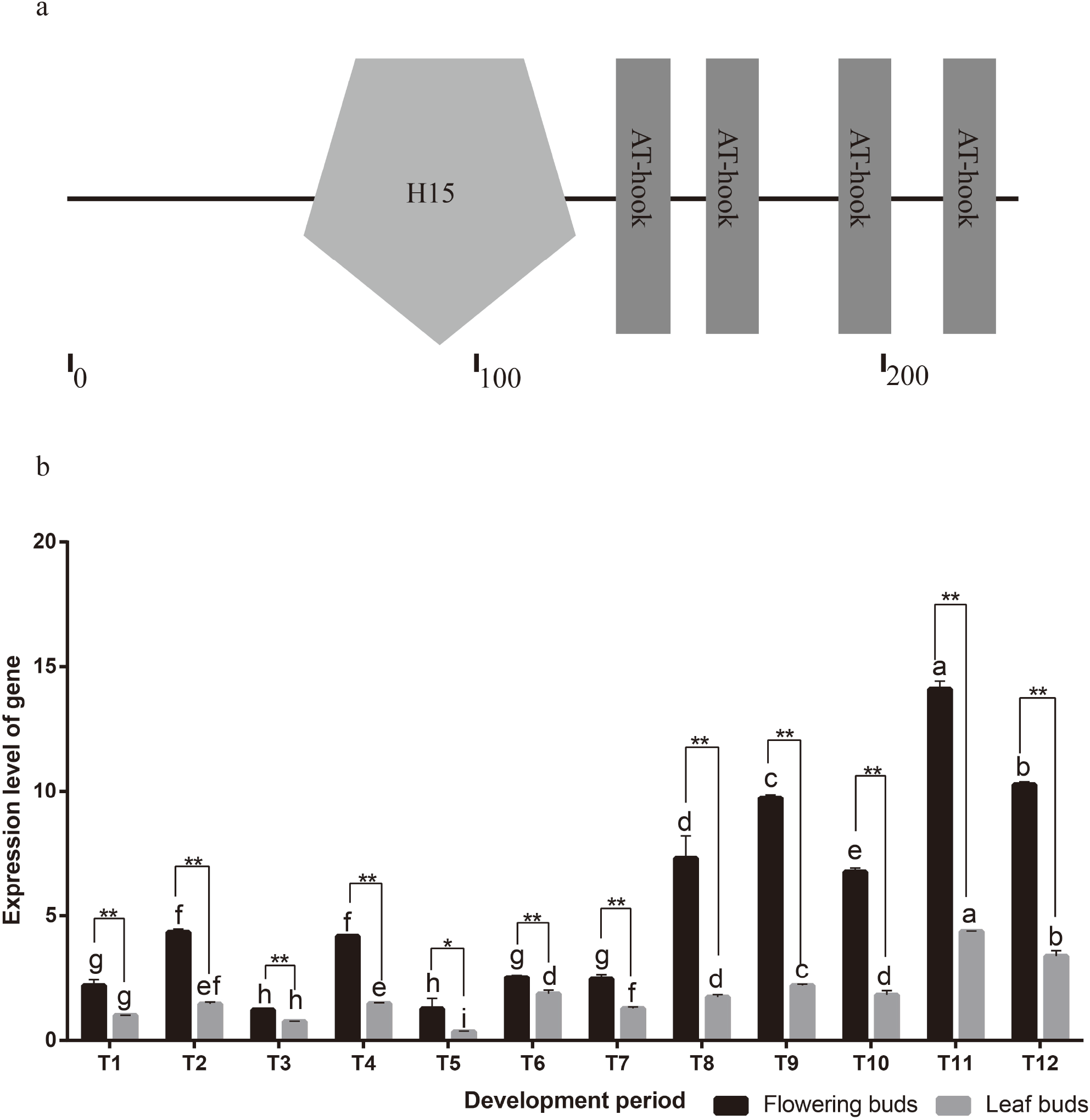
The sequence and expression profile analysis of *CbuHMGA*. a) Sequence analysis of the CbuHMGA protein by SMART. b) The expression analysis of *CbuHMGA* between flowering buds and leaf buds during the development periods. T1-T3 represents the flowering buds and leaf buds collected in the dormant period; T4-T5 represents the flowering buds and leaf buds collected in the germination period; T6-T9 represents the flowering buds and leaf buds collected in the floral transition period; and T10-T12 represents the flowering buds and leaf buds collected in the reproductive period. Notably, even though floral transition was never observed in the leaf buds, the leaf buds collected in the period corresponding to floral transition are henceforth called the floral transition period and reproductive period for convenience. Error bars indicate SD from three independent experiments. *Difference between flowering buds (black) and leaf buds (gray) is significant (Student’s test; p<0.05). **Difference between flowering buds (black) and leaf buds (gray) is highly significant (Student’s test; p<0.01).

### Localization of the CbuSPL9 and CbuHMGA proteins

We fused GFP at the C-terminus of CbuSPL9 and CbuHMGA and transformed them into *Nicotiana benthamiana* leaf epidermal cells to determine the localization of the CbuSPL9 and CbuHMGA proteins. 35S:GFP was used as a control. The GFP fluorescence in CbuSPL9-GFP and CbuHMGA-GFP was exclusively observed in the nucleus, whereas the fluorescence of GFP in the control was distributed throughout the entire cell (Figure 5a). These results showed that CbuSPL9 and CbuHMGA were located in the nucleus. This result was in agreement with the prediction based on the protein structure.

**Figure 5.**
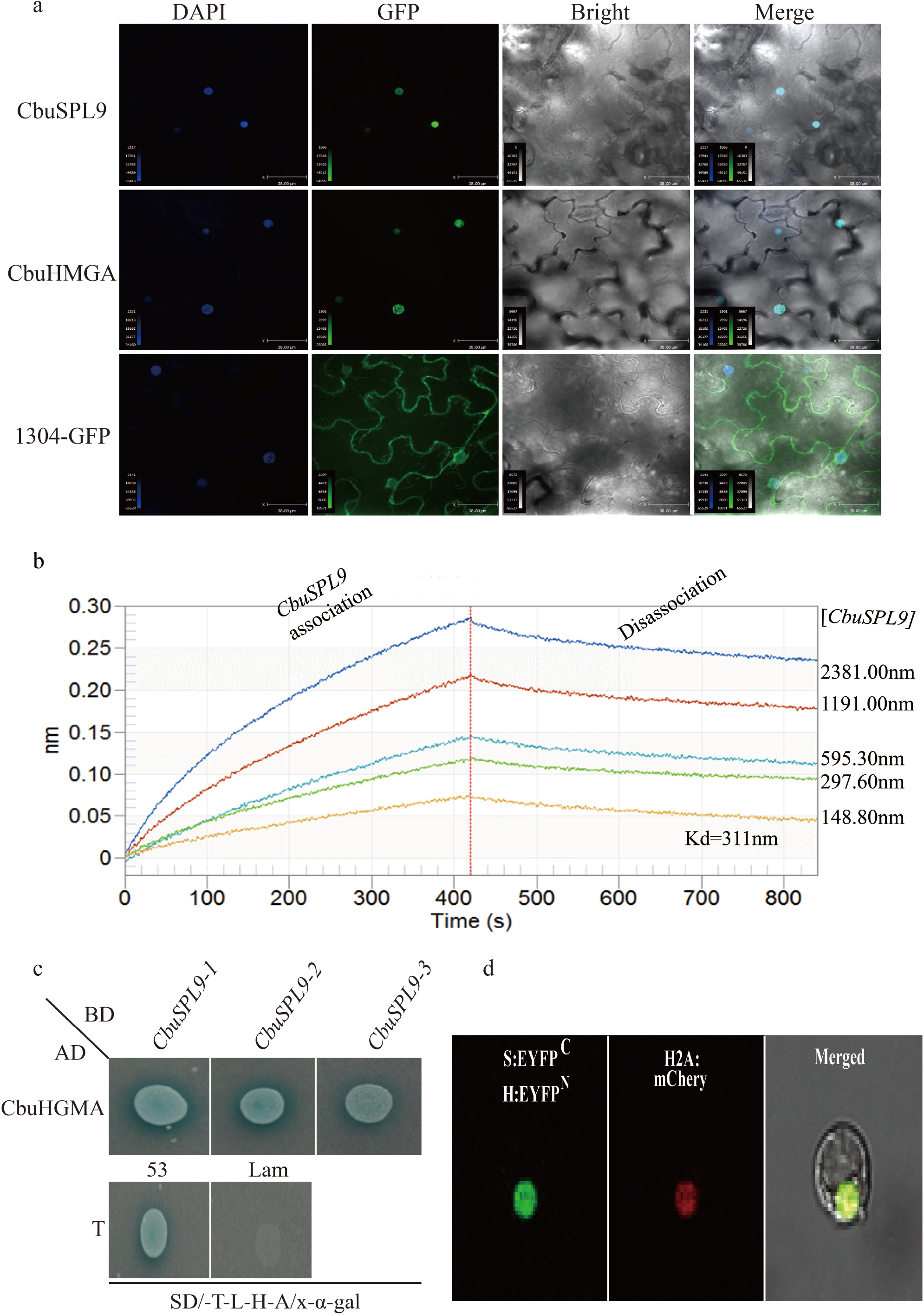
Interaction between the two proteins CbuSPL9 and CbuHMGA. a) Nuclear localization of the CbuSPL9 protein and CbuHMGA protein. The GFP (control) gene, CbuSPL9-GFP fusion gene and CbuHMGA-GFP fusion gene were expressed transiently in *Nicotiana benthamiana* leaf epidermal cells and observed with confocal microscopy. DAPI, DAPI for nuclear staining image; GFP, GFP green fluorescence image; Merge, the merged images of bright-field, GFP and DAPI staining. b) Binding of GST-CbuSPL9 to HIS-CbuHMGA, with GST-CbuSPL9 concentrations of 2381.00 nm, 1191.0 nm, 595.30 nm, 297.60 nm, 148.80 nm assessed by real-time biolayer interferometry. c) Yeast two-hybrid assays for the interactions between CbuSPL9 and CbuHMGA. CbuHMGA (as prey) was fused with the GAL4 activation domain (AD) in pGADT7, while CbuSPL9 (as bait) was fused with the GAL DNA-binding domain (BD) in pGBKT7. The positive control was as follows: pGADT7-T and pGBK-53. The negative control was as follows: pGADT7-T and pGBK-Lam. Interactions are indicated by the blue color on SD/-Trp/-Leu/-His/-Ade/X-α-gal medium. d) Confocal images of the BiFC analysis in protoplasts from *P. trichocarpa*. CbuSPL9 was fused with EYFP^C^, and CbuHMGA was fused with EYFP^N^. EYFP signal was detected in the nucleus of the S1-21 protoplasts transfected by H2A:mCherry and CbuSPL9:EYFP^C^ with (A) CbuHMGA:EYFP^N^. Bars=50 µm.

### Protein interaction analysis

Biolayer interferometry (BLI) was used to determine dissociation constants (K_D_) as well as the on-and off-rate (k_on_ and k_off_) for HIS-CbuHMGA binding to GST-CbuSPL9 (Table 2). Five different concentrations of GST-CbuSPL9 (2381.00 nm, 1191.00 nm, 595.30 nm, 297.60 and 148.80 nm) were evaluated, and the K_D_ was 311 nm for each tested concentration. These results suggested a strong interaction between CbuSPL9 and CbuHMGA (Figure 5b).

**Table 2.** Interaction between HIS-CbuHMGA and GST-CbuSPL9 by BLI

| Protein1 | Protein2 | Conc. of Protein2/nM | $K_{on}/Ms$ | $K_{off}/s$ | $K_D/nM$ |
| --- | --- | --- | --- | --- | --- |
| HIS-CbuHMGA | GST-CbuSPL9 | 2381 | 5.40E-04 | 1.74E+03 | 311 |
| HIS-CbuHMGA | GST-CbuSPL9 | 1191 | 5.40E-04 | 1.74E+03 | 311 |
| HIS-CbuHMGA | GST-CbuSPL9 | 595.3 | 5.40E-04 | 1.74E+03 | 311 |
| HIS-CbuHMGA | GST-CbuSPL9 | 297.6 | 5.40E-04 | 1.74E+03 | 311 |
| HIS-CbuHMGA | GST-CbuSPL9 | 148.8 | 5.40E-04 | 1.74E+03 | 311 |

To confirm these interactions, the full-length cDNA of *CbuSPL9* was inserted into the vector pGBKT7 (BD-CbuSPL9) as bait, and the full-length cDNA of *CbuHMGA* was inserted into the vector pGADT7 (AD-CbuHMGA) as prey. Yeast strains containing AD-CbuHMGA and BD-CbuSPL9 were positive for X-α-gal activity when grown on synthetically defined (SD)/-Trp/-Leu/-His/-Ade medium (Figure 5c). These results showed that CbuSPL9 interacted with HMGA in yeast. Finally, a bimolecular fluorescence complementation (BiFC) assay was performed. CbuSPL:EYFP^C^ was cotransfected with CbuHMGA:EYFP^N^ and H2A:mCherry into protoplasts. The signal of enhanced yellow fluorescent protein (EYFP) was colocalized with mCherry (Figure 5d). Collectively, we demonstrated that CbuSPL9 formed a heterodimer with CbuHMGA in the nucleus.

### The overexpression of *CbuHMGA* in Arabidopsis

To characterize the potential function of *CbuHMGA*, we overexpressed *CbuHMGA* in Arabidopsis. The *oe-hmga* transgenic lines developed aberrant flowers with abnormal petals and stamens (Figure 6aI-VI). This mutant phenotype was similar to the phenotype of floral organs when *CbuSPL9* was overexpressed (Supplementary Table S5). However, flowering time was not affected in the *oe-hmga* transgenic lines compared to that in the col line (Supplementary Table S6). The expression of endogenous *AtSPL9* was thoroughly studied to further monitor flower development in the transgenic lines. Within the sampling period, the expression level of *AtSPL9* continuously increased in *oe-hmga* and wild type. However, in *oe-spl9*, the expression of *AtSPL9* reached a peak in T6 and then decreased (Figure 6b). The shift in *AtSPL9* expression in *oe-spl9*, but not in *oe-hmga*, further confirmed the observation that *CbuHMGA* cannot accelerate flower development, while *CbuSPL9* can accelerate flower development.

**Figure 6.**
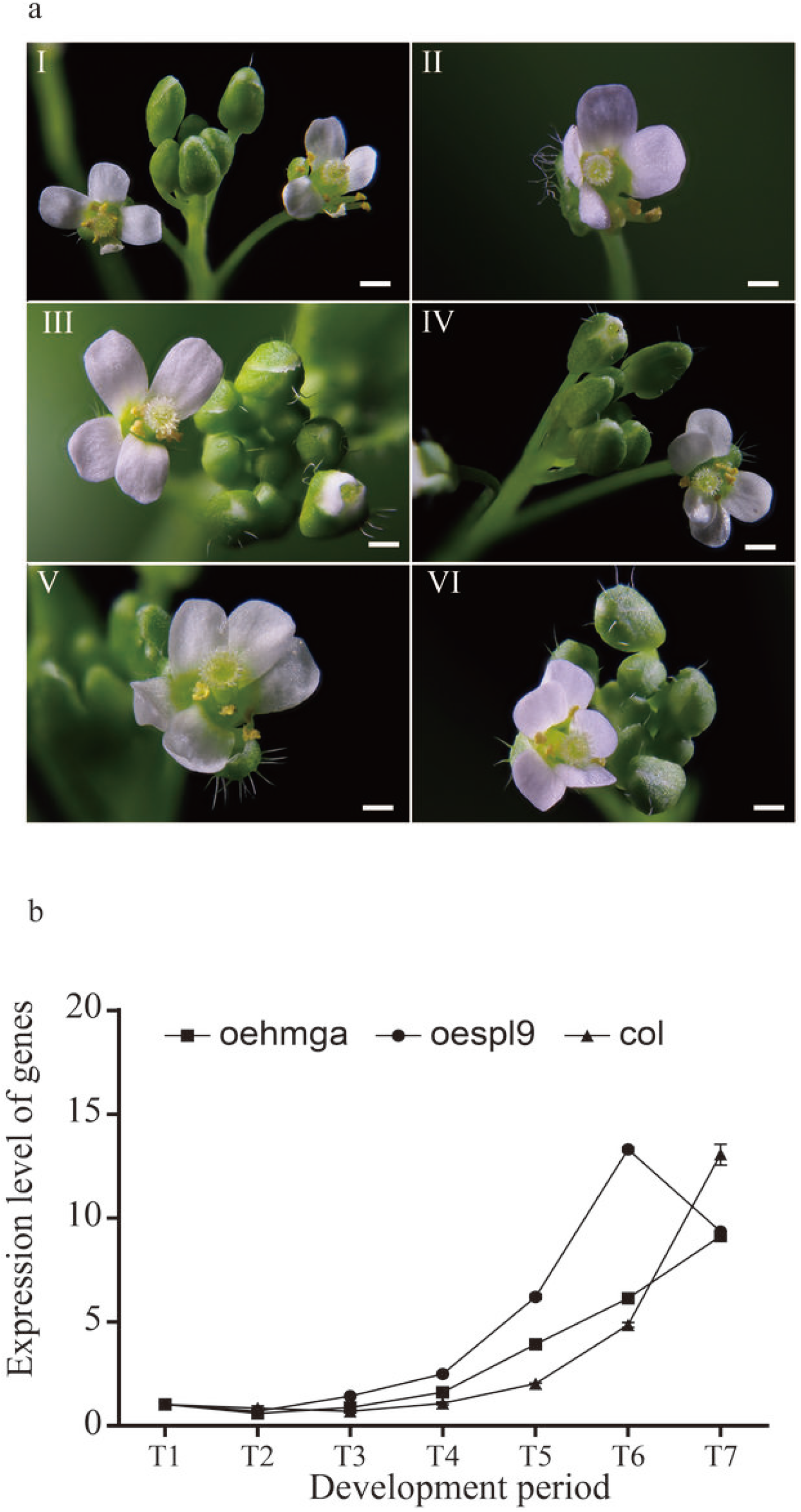
Phenotype of overexpression mutants of *CbuHMGA*. a) The change in floral organs occurred in the overexpression of the CbuHMGA transgenic Arabidopsis lines. The images are the floral organ mutant from oe-hmga transgenic Arabidopsis lines (a/I-VI). b) Expression profiles of atSPL9 homologous genes in the oespl9 transgenic plants, oe-hmga transgenic plants and col. represents the PCR results in oe-hmga; represents the PCR results in oespl9; and represents the PCR results in col. Three independent biological replicates were performed, and each replicate was measured in triplicate. The error bars show the standard deviation of the results of three technical replicates.

## Discussion

Flowering is a very complex process, a qualitative change in the life history of higher plants, and a central link in plant development. “Bairihua”, a variety of the flowering perennial woody plant *C. bungei*, has a large amount of flowers and a long flowering period. “Bairihua” is an excellent material for evaluating the flowering process in trees.

*SPL*s play an important role in regulating flowering in many plants, most notably in Arabidopsis. In woody plants, *SPLs* are studied extensively, but limited information about flowering is known^29, 48–51^. Here, we cloned the SBP-domain-encoding gene *CbuSPL9* in *C. bungei.* Its conserved structure and flower development-related expression trends suggest that *CbuSPL9* might have conserved *SPL9* functions in *C. bungei.* Furthermore, heterogeneous overexpression of *CbuSPL9* in Arabidopsis accelerated flower development and lead to aberrant flower organs. Although we could not generate homogenous transgenic plants due to technology and time, these results are sufficient to demonstrate that *CbuSPL9* is an SPL homolog and participates in flower development.

TFs can affect biological processes by forming complexes with other TFs. However, most of the studies on SPLs have focused on their interactions with DNA motifs^38–40^. The proteins that could directly interact with SPLs and participate in flower regulation are largely unknown. For further insight, we screened the yeast hybridization library for CbuSPL9 in buds. Many flowering-related proteins were detected, for example, the GATA-related proteins^52–57^, bHLH48^58^, FLA^59–62^, and PHD finger protein ALFIN-LIKE^63–65^. Furthermore, a high mobility group (HMG) protein, CbuHMGA, was identified to interact directly with CbuSPL9. ChuHMGA contains an H15 domain and 4 AT-hook motifs. The BiFC assay and BLI assay further confirmed their interaction. The BiFC assay suggested that CbuSPL9 and CbuHMGA could form protein complexes in the nucleus. HMGA proteins have a highly conserved structure^66, 67^. Unlike the HMGAs in the animal kingdom, which invariably contain three AT-hook motifs, most plant HMGA proteins have four AT-hook motifs^67, 68^. Additionally, the amino-terminal region of plant HMGA proteins shares remarkable homology with the DNA-binding domain of histone H1^69, 70^. A similar structural description was confirmed in CbuHMGA, which encoded a H15 domain and four AT-hook motifs. HMGA is an architectural TF^70–73^. It regulates gene expression in vivo by controlling the formation of multiprotein complexes on the AT-rich regions of certain gene promoters^68, 70, 73–80, 81^. To date, some plant HMGA proteins have been isolated, such as those in soybean^82^, rice^83^, maize^84^ and Arabidopsis^67, 73, 75, 80, 85–87^. The Arabidopsis HMGA gene was detected in flowers, and developing siliques had the highest expression^87^. However, studies of the interacting proteins and the function of HMGA in woody plants are rare. In “Bairihua”, the expression of *CbuHMGA* was elevated in the dormant period, and the highest expression level was detected in the reproductive growth period. The result was consistent with that in Arabidopsis. Although the *oe-hmga* transgenic lines did not show a flowering time phenotype, the floral organ mutation of *oe-hmga* was similar to that of the *oe-spl9* transgenic lines.

The HMGA proteins have an important role in biological processes and interact with different TFs^70, 73, 88^. Their intrinsic flexibility allows the HMGA proteins to participate in specific protein-DNA and protein-protein interactions that induce structural changes in chromatin and the formation of stereospecific complexes called ‘enhanceosomes’ on the promoter/enhancer regions of genes whose transcription they regulate. The chromatin structure changes affect the ability of TFs to bind with the promoter/enhancer regions^68, 74, 77, 78, 80, 85^. From the results in this study, we suggest that the interaction of CbuHMGA with CbuSPL9 might strengthen or weaken the binding ability of CbuSPL9 with the corresponding DNA sequences or downstream proteins, thus affecting the flowering process. Further experiments are needed to test this hypothesis.

## Conclusion

“Bairihua” provides us with a valuable opportunity to gain a deeper understanding of the flowering process of woody plants. Our study showed that *SPLs* may have similar structures and regulatory mechanisms in perennial trees and Arabidopsis. Screening CbuSPL9-interacting proteins revealed not only additional proteins for research on the regulatory pathway of CbuSPL9 but also the CbuSPL9 interaction with CbuHMGA. *oe-hmga* showed a phenotype that affected floral organ development but did not change flowering time. This result suggests that the mechanism by which *CbuSPL9* affects the flowering process is a complex process, and *CbuHMGA* is involved in the development of flower organs but not in the regulation of flowering time.

## Author contributions

J.H.W., S.G.Z., G.Z.Q., L.I.S., Z.W. and W.J.M. designed the experiments. Z.W. and W.J.M. analyzed the RNA-seq data and wrote the manuscript. Z.W., and T.Q.Z. detected the expression of genes using qRT-PCR. Z.W., N.L., F.Q.O.Y., N.W. and G.J.Y. collected the samples used in the experiment. All the authors have read the paper and agreed to list their names as co-authors.

## Supplementart Data

**Supplementary Figure S1.**
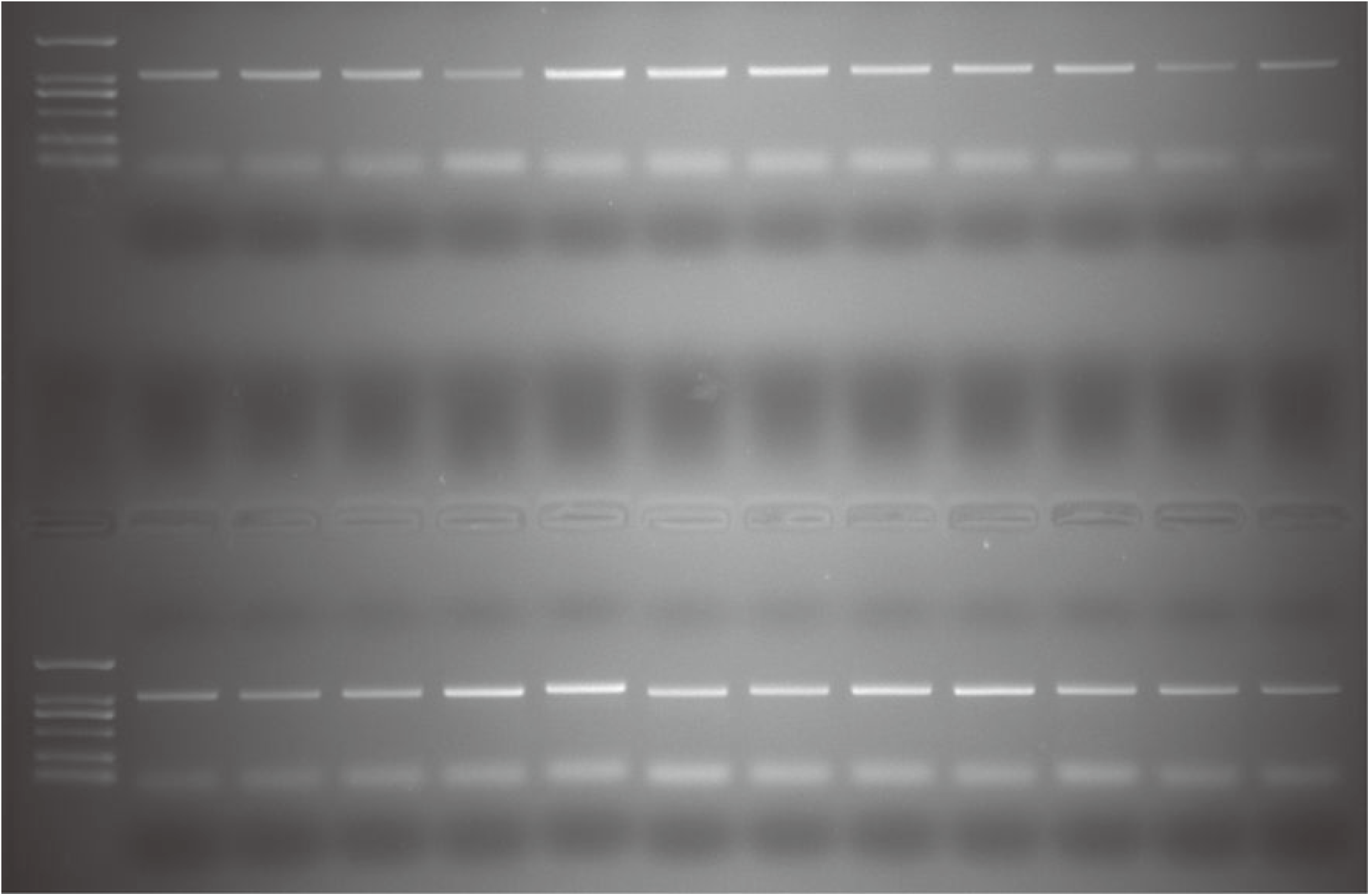
Verification of *oe-SPL9* positive Arabidopsis.

**Supplementary Figure S2.**
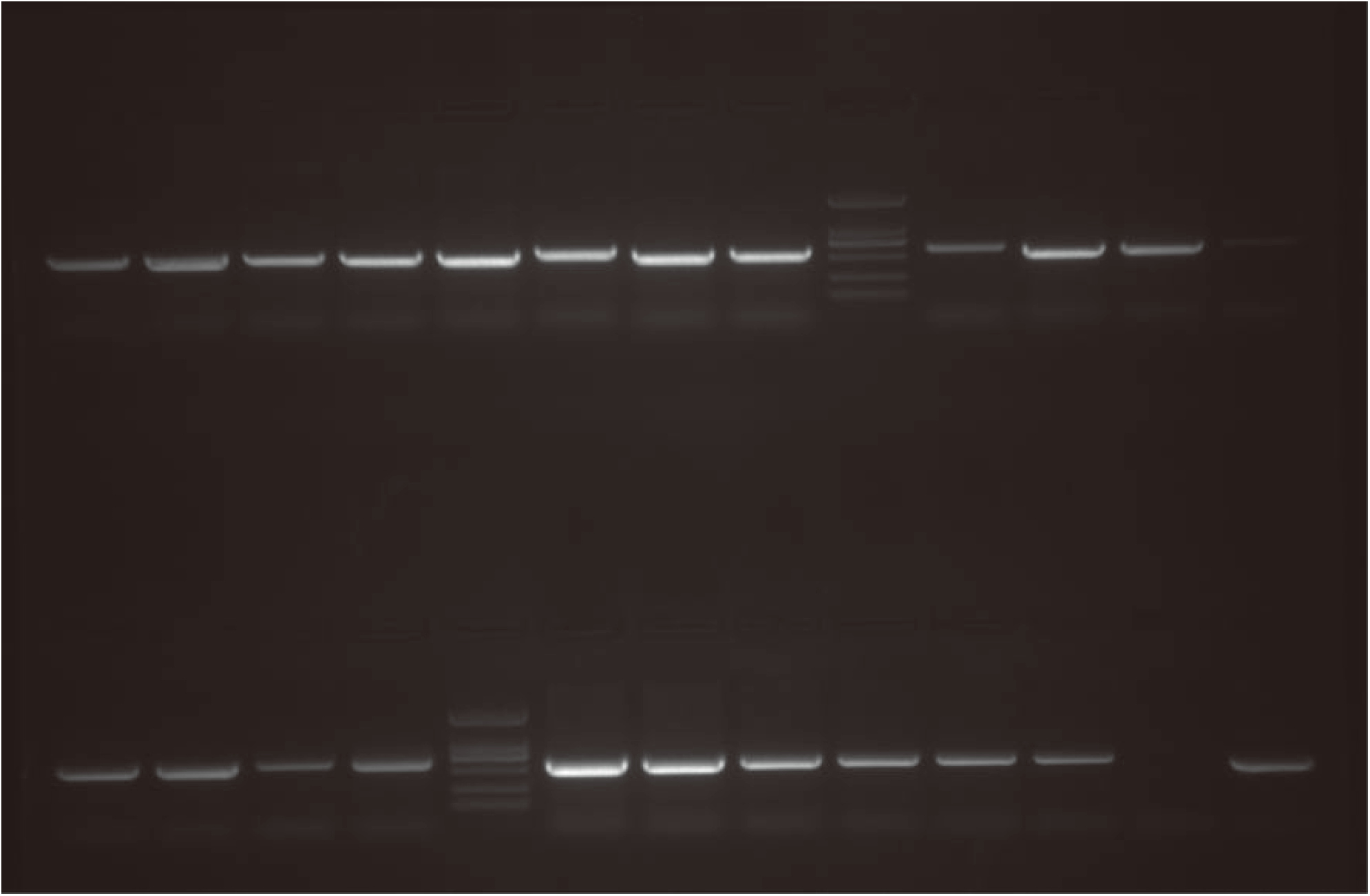
Verification of *oe-HMGA* positive Arabidopsis.

**Supplementary Table S1.**
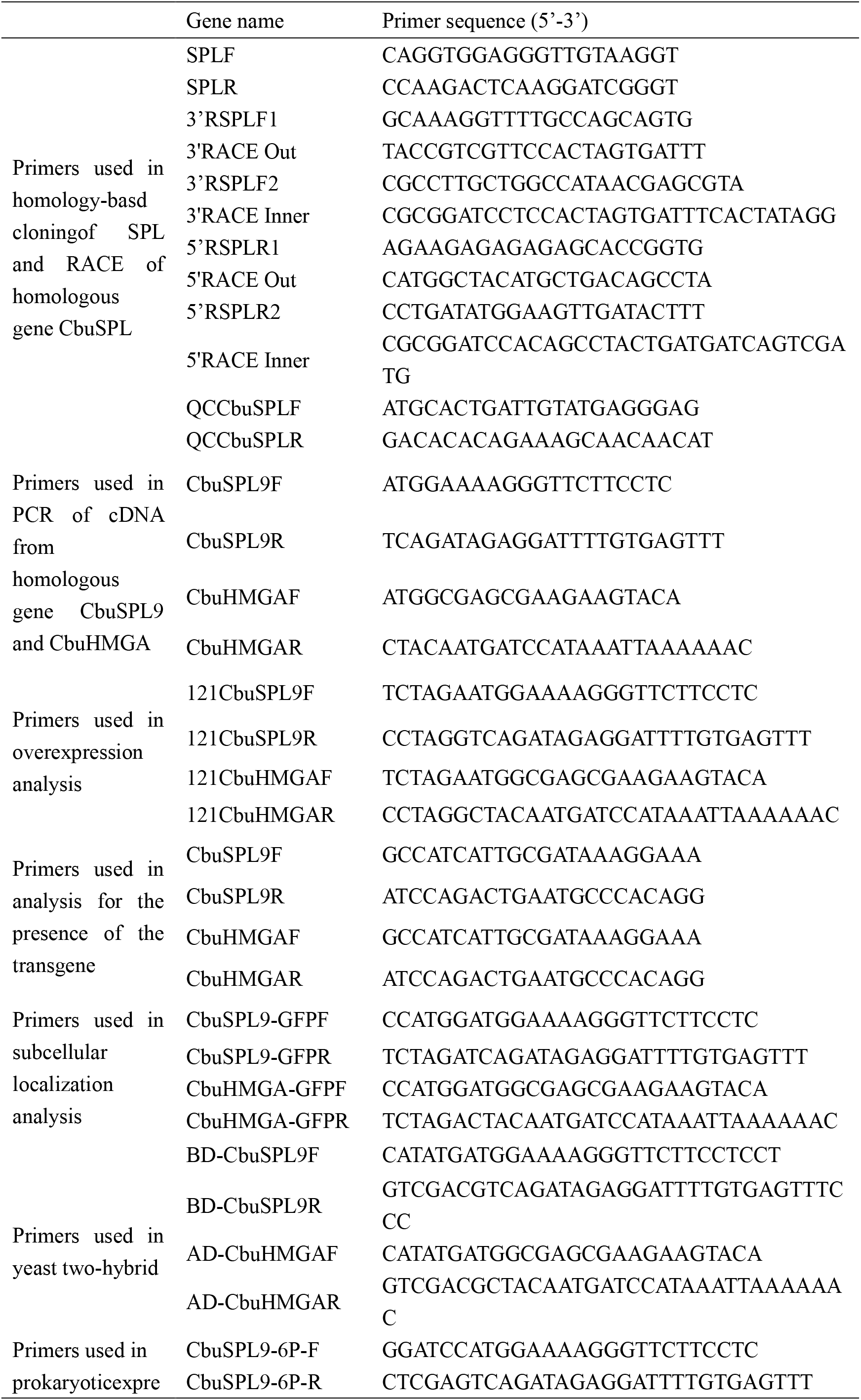
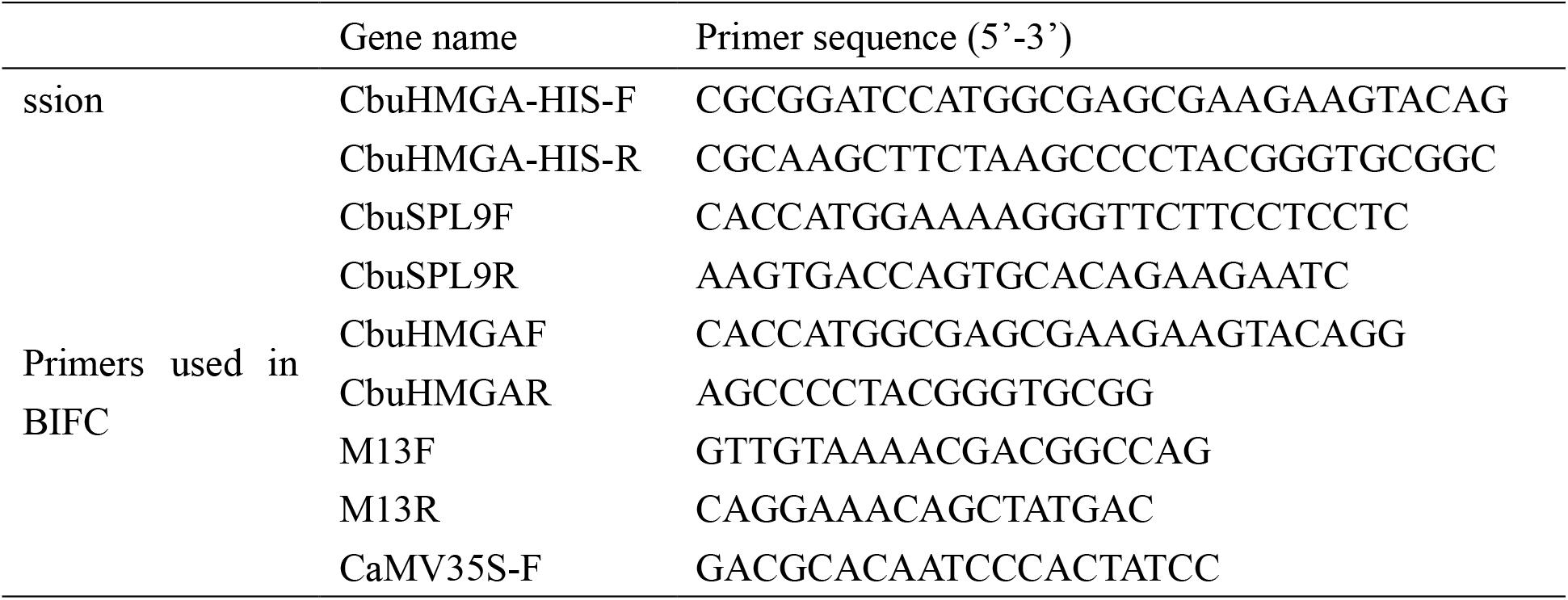
The list of the all Primers used in this paper.

**Supplementary Table S2.**
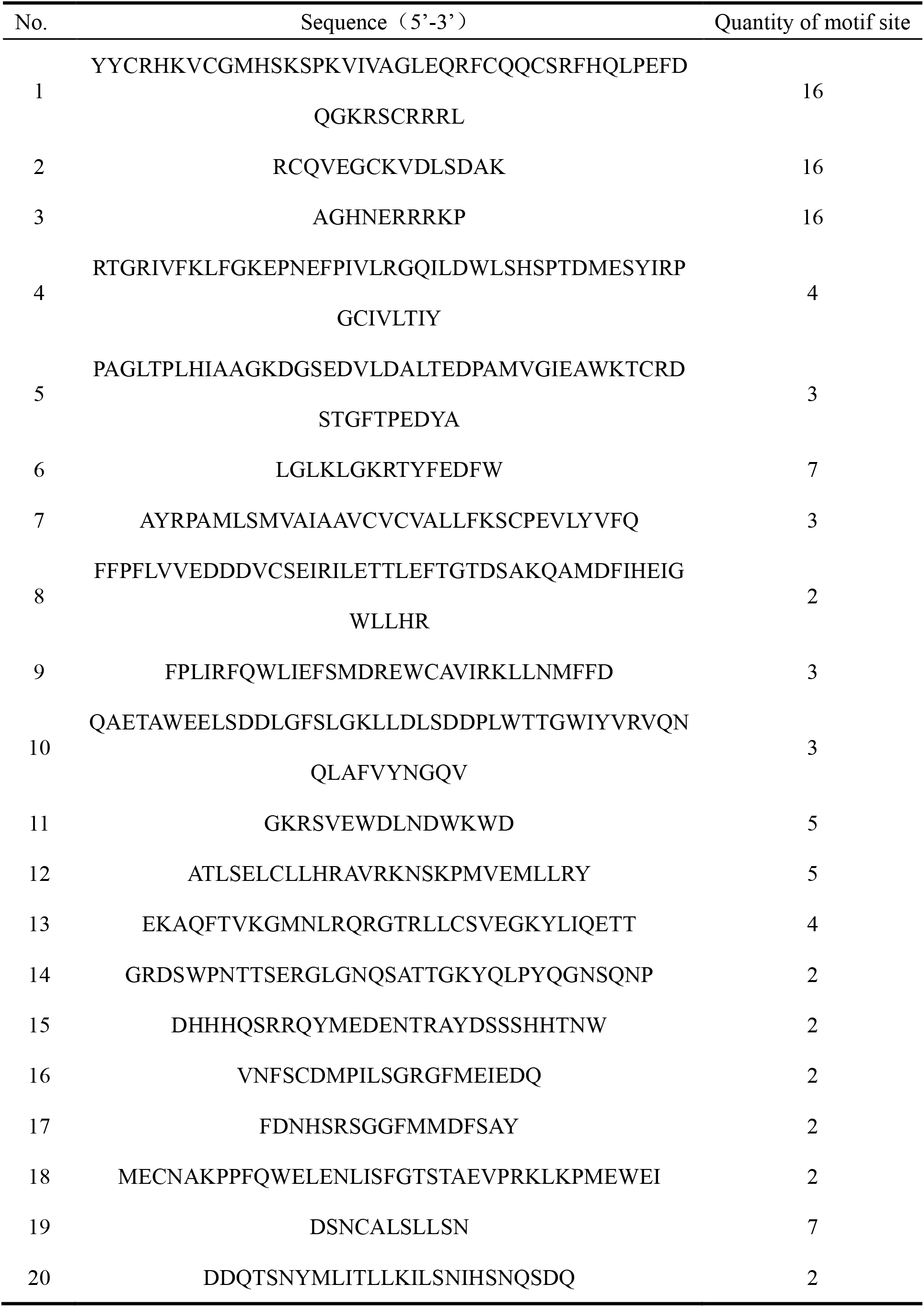
Details of motif-sequences of CbuSPL9 were identified by MEME.

**Supplementary Table S3.**
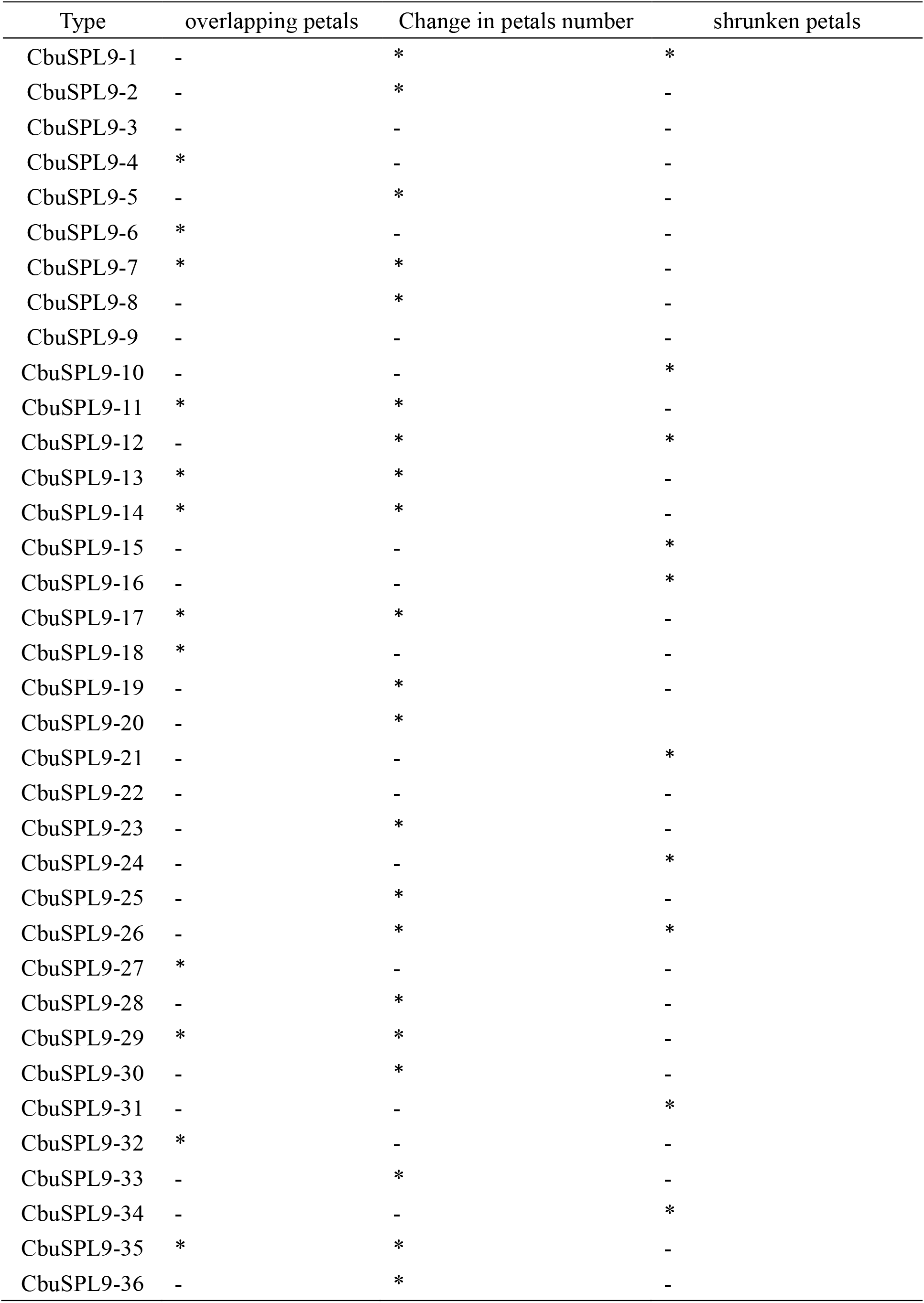
Statistics of mutant of floral organs in *oe-SPL9* Arabidopsis.

**Supplementary Table S4.** Statistics of mutant of flowering time in *oe-SPL9* Arabidopsis.

| Type | DAY | Type | DAY |
| --- | --- | --- | --- |
| CbuSPL9-01 | 22 | WT-01 | 27 |
| CbuSPL9-02 | 17 | WT-02 | 27 |
| CbuSPL9-03 | 23 | WT-03 | 26 |
| CbuSPL9-04 | 21 | WT-04 | 27 |
| CbuSPL9-05 | 22 | WT-05 | 25 |
| CbuSPL9-06 | 16 | WT-06 | 28 |
| CbuSPL9-07 | 22 | WT-07 | 28 |
| CbuSPL9-08 | 23 | WT-08 | 26 |
| CbuSPL9-09 | 23 | WT-09 | 25 |
| CbuSPL9-10 | 22 | WT-10 | 27 |
| CbuSPL9-11 | 21 | WT-11 | 25 |
| CbuSPL9-12 | 22 | WT-12 | 27 |
| CbuSPL9-13 | 23 | WT-13 | 27 |
| CbuSPL9-14 | 22 | WT-14 | 26 |
| CbuSPL9-15 | 22 | WT-15 | 27 |
| CbuSPL9-16 | 24 | WT-16 | 27 |
| CbuSPL9-17 | 22 | WT-17 | 25 |
| CbuSPL9-18 | 18 | WT-18 | 29 |
| CbuSPL9-19 | 23 | WT-19 | 27 |
| CbuSPL9-20 | 21 | WT-20 | 28 |
| CbuSPL9-21 | 24 | WT-21 | 27 |
| CbuSPL9-22 | 23 | WT-22 | 26 |
| CbuSPL9-23 | 21 | WT-23 | 26 |
| CbuSPL9-24 | 24 | WT-24 | 27 |
| CbuSPL9-25 | 21 | WT-25 | 27 |
| CbuSPL9-26 | 23 | WT-26 | 25 |
| CbuSPL9-27 | 21 | WT-27 | 26 |
| CbuSPL9-28 | 18 | WT-28 | 26 |
| CbuSPL9-29 | 21 | WT-29 | 27 |
| CbuSPL9-30 | 20 | WT-30 | 26 |

**Supplementary Table S5.** Statistics of mutant of floral organs in *oe-HMGA* Arabidopsis.

| Type | Overlapping petals | Change in petals number | shrunk petals |
| --- | --- | --- | --- |
| CbuHMGA-1 | * | * |  |
| CbuHMGA-2 |  |  | * |
| CbuHMGA-3 |  | * |  |
| CbuHMGA-4 | - | - | - |
| CbuHMGA-5 |  | * | * |
| CbuHMGA-6 |  | * |  |
| CbuHMGA-7 | * | * |  |
| CbuHMGA-8 |  | * |  |
| CbuHMGA-9 |  |  | * |
| CbuHMGA-10 | * | * |  |
| CbuHMGA-11 | - | - | - |
| CbuHMGA-12 | * | * |  |
| CbuHMGA-13 |  | * |  |
| CbuHMGA-14 |  | * |  |
| CbuHMGA-15 |  |  | * |
| CbuHMGA-16 |  | * |  |
| CbuHMGA-17 |  | * | * |
| CbuHMGA-18 |  | * |  |
| CbuHMGA-19 | - | - | - |
| CbuHMGA-20 | - | - | - |
| CbuHMGA-21 |  | * |  |
| CbuHMGA-22 | * |  |  |
| CbuHMGA-23 |  |  | * |
| CbuHMGA-24 |  | * |  |
| CbuHMGA-25 |  |  | * |
| CbuHMGA-26 | * | * |  |
| CbuHMGA-27 |  |  | * |
| CbuHMGA-28 | - | - | - |
| CbuHMGA-29 |  |  | * |
| CbuHMGA-30 | * | * |  |
| CbuHMGA-31 |  |  | * |
| CbuHMGA-32 |  | * |  |
| CbuHMGA-33 |  |  | * |
| CbuHMGA-34 |  | * |  |
| CbuHMGA-35 | - | - | - |
| CbuHMGA-36 | * | * |  |

**Supplementary Table S6.** Statistics of mutant of flowering time in *oe-HMGA* Arabidopsis.

| Type | Time | Type | Time |
| --- | --- | --- | --- |
| CbuHMGA-01 | 25 | WT-01 | 27 |
| CbuHMGA-02 | 25 | WT-02 | 27 |
| CbuHMGA-03 | 26 | WT-03 | 26 |
| CbuHMGA-04 | 24 | WT-04 | 27 |
| CbuHMGA-05 | 27 | WT-05 | 25 |
| CbuHMGA-06 | 28 | WT-06 | 28 |
| CbuHMGA-07 | 26 | WT-07 | 28 |
| CbuHMGA-08 | 24 | WT-08 | 26 |
| CbuHMGA-09 | 28 | WT-09 | 25 |
| CbuHMGA-10 | 27 | WT-10 | 27 |
| CbuHMGA-11 | 24 | WT-11 | 25 |
| CbuHMGA-12 | 28 | WT-12 | 27 |
| CbuHMGA-13 | 27 | WT-13 | 27 |
| CbuHMGA-14 | 26 | WT-14 | 26 |
| CbuHMGA-15 | 25 | WT-15 | 27 |
| CbuHMGA-16 | 28 | WT-16 | 27 |
| CbuHMGA-17 | 25 | WT-17 | 25 |
| CbuHMGA-18 | 24 | WT-18 | 29 |
| CbuHMGA-19 | 28 | WT-19 | 27 |
| CbuHMGA-20 | 27 | WT-20 | 28 |
| CbuHMGA-21 | 26 | WT-21 | 27 |
| CbuHMGA-22 | 26 | WT-22 | 26 |
| CbuHMGA-23 | 26 | WT-23 | 26 |
| CbuHMGA-24 | 25 | WT-24 | 27 |
| CbuHMGA-25 | 27 | WT-25 | 27 |
| CbuHMGA-26 | 25 | WT-26 | 25 |
| CbuHMGA-27 | 27 | WT-27 | 26 |
| CbuHMGA-28 | 26 | WT-28 | 26 |
| CbuHMGA-29 | 26 | WT-29 | 27 |
| CbuHMGA-30 | 28 | WT-30 | 26 |

